# Effects of CO_2_ and RuBisCO concentration on cyanobacterial growth and carbon isotope fractionation

**DOI:** 10.1101/2021.04.20.440233

**Authors:** Amanda Garcia, Mateusz Kedzior, Arnaud Taton, Meng Li, Jodi Young, Betül Kaçar

## Abstract

Carbon isotope biosignatures preserved in the Precambrian geologic record are primarily interpreted to reflect ancient cyanobacterial carbon fixation catalyzed by Form I RuBisCO enzymes. The average range of isotopic biosignatures generally follows that produced by extant cyanobacteria. However, this observation is difficult to reconcile with several environmental (e.g., temperature, pH, and CO_2_ concentrations), molecular, and physiological factors that likely would have differed during the Precambrian and can produce fractionation variability in contemporary organisms that meets or exceeds that observed in the geologic record. To test a range of genetic and environmental factors that may have impacted ancient carbon isotope biosignatures, we engineered a mutant strain of the model cyanobacterium *Synechococcus elongatus* PCC 7942 that overexpresses RuBisCO and characterized the resultant physiological and isotope fractionation effects. We specifically investigated how both increased atmospheric CO_2_ concentrations and RuBisCO regulation influence cell growth, oxygen evolution rate, and carbon isotope fractionation in cyanobacteria. We found that elevated CO_2_ increases the growth rate of wild-type and mutant strains, and that the pool of active RuBisCO enzyme increases with increased expression. RuBisCO overexpression in our engineered strain does not significantly affect isotopic discrimination at all tested CO_2_ concentrations, yielding cellular ^13^C/^12^C isotope discrimination (ε_p_) of ∼24‰ for both wild-type and mutant strains at elevated CO_2_. Understanding the environmental factors that impact gene regulation, physiology, and evolution is crucial for reconciling microbially driven carbon isotope fractionation with the geologic record carbon biosignatures.

## INTRODUCTION

The conserved microbial metabolic pathways that drive global biogeochemistry emerged on Earth billions of years ago, the evolution of which has both shaped and been shaped by large-scale environmental transitions (Falkowski, Fenchel, & Delong, 2008; Knoll & Nowak, 2017; Lyons, Fike, & Zerkle, 2015). These microbial processes have left distinct signatures that are evidence of biological activity billions of years in the past. The oldest and most extensive signature of biological activity on Earth is the deviation in stable carbon isotopic compositions (^13^C/^12^C, expressed as δ^13^C) between preserved inorganic and organic carbon, interpreted to reflect the isotopic discrimination of ancient biological carbon fixation (Des Marais, 2001; Krissansen-Totton, Buick, & Catling, 2015; Lloyd et al., 2020; Schidlowski, 2001). This deviation is primarily shaped by enzymes that preferentially assimilate the lighter ^12^C isotope from inorganic carbon sources. Carbon biosignatures preserved in the geologic record therefore reflect the long-term evolution of these enzyme-mediated processes and their hosts’ physiologies.

The RuBisCO enzyme (ribulose 1,5-bisphosphate (RuBP) carboxylase/oxygenase) catalyzes the reduction of inorganic CO_2_ as the initial step of carbon assimilation into organic biomass via the Calvin-Benson-Bassham (CBB) cycle (Erb & Zarzycki, 2018; Nisbet et al., 2007; Tcherkez, Farquhar, & Andrews, 2006). RuBisCO is one of the most abundant proteins on Earth (Bar-On & Milo, 2019; Ellis, 1979; Raven, 2013), and its presence in photoautotrophic organisms, including early-evolved cyanobacteria, suggests that this enzyme has played a significant role in primary production for much of Earth history (Hamilton, Bryant, & Macalady, 2016; Schirrmeister, Sanchez-Baracaldo, & Wacey, 2016; Schopf, 2011). Thus, the isotopic fractionation behavior of RuBisCO is also thought to have constrained Precambrian carbon isotope signatures preserved in the geologic record (Garcia, Cavanaugh, & Kacar, 2021; Schidlowski, 1988). Though there are multiple forms of RuBisCO, the known range of the Form I RuBisCO isotopic effect (ε ≈ 20-30‰) (Guy, Fogel, & Berry, 1993; Scott et al., 2007; von Caemmerer, Tazoe, Evans, & Whitney, 2014) — the form utilized by extant cyanobacteria and responsible for the bulk of modern primary production (Field, 1998) — is largely consistent with the ∼25‰ mean deviation between preserved inorganic and organic carbon isotopic compositions across geologic time (Des Marais, 2001; Havig, Hamilton, Bachan, & Kump, 2017; Kedzior et al., 2022; Krissansen-Totton et al., 2015; Lloyd et al., 2020; Schidlowski, 2001). The isotopic effects of cyanobacterial Form IA and IB RuBisCO in particular have been measured between ∼22 to 24‰ (Guy et al., 1993; Scott et al., 2007), whereas land plant Form IB RuBisCO typically has larger effects of up to ∼29‰ (von Caemmerer et al., 2014) and Form ID RuBisCO have exhibited effects as small as ∼11‰ (Boller, Thomas, Cavanaugh, & Scott, 2011).

RuBisCO is an important factor in the generation of distinct carbon isotopic biosignatures associated with CBB-utilizing organisms (Hayes, 2001; Laws, Popp, Bidigare, Kennicutt, & Macko, 1995; Wilkes & Pearson, 2019). Nonetheless the carbon isotopic composition of bulk photoautotrophic biomass often deviates from values obtained for purified RuBisCO, a discrepancy at least partially explained by the fact that the latter are measured at CO_2_ saturation (e.g., (Boller et al., 2011; Guy et al., 1993; Scott et al., 2007; von Caemmerer et al., 2014)). This discrepancy might also be attributable to several intracellular physiological and metabolic features that additionally shift the isotopic composition in organism biomass. These include the activation of carbon-concentrating mechanisms (CCM) that elevate CO_2_ in RuBisCO-containing cellular compartments (Hurley, Wing, Jasper, Hill, & Cameron, 2021; Laws, Popp, Cassar, & Tanimoto, 2002; Price, Badger, Woodger, & Long, 2008; Raven, Cockell, & De La Rocha, 2008; Wilkes & Pearson, 2019) and the diffusive transport of CO_2_ (Hayes, 1993; Rau, Riebesell, & Wolf-Gladrow, 1996). Studies have shown that photosynthetic carbon isotope discrimination (ε_p_) may vary due to environmental factors and cellular physiological responses, including temperature (Deleens, Treichel, & O’Leary, 1985; Wong & Sackett, 1978), pH (Hinga, Arthur, Pilson, & Whitaker, 1994; Roeske & O’Leary, 1984), growth rate (Bidigare et al., 1997; Laws, Bidigare, & Popp, 1997), and CO_2_ concentration (Eichner, Thoms, Kranz, & Rost, 2015; Freeman & Hayes, 1992; Hinga et al., 1994; Hurley et al., 2021; Schubert & Jahren, 2012; Wilkes, Lee, McClelland, Rickaby, & Pearson, 2018). These variations indicate a compelling need to comprehensively characterize both internal and external factors that can affect isotope fractionation, though the study of each tend to be siloed in biological and geobiological fields, respectively.

An aspect of photoautotrophic carbon isotopic discrimination that has not been thoroughly investigated is how RuBisCO expression levels may have influenced ancient host organism growth and biosignature generation when CO_2_ leves were higher. RuBisCO expression has been shown to be CO_2_ -sensitive (Gesch et al., 2003; Onizuka et al., 2002; Sengupta, Sunder, Sohoni, & Wangikar, 2019). This observation is particularly important, considering that atmospheric CO_2_ concentrations exceeded present day levels by more than an order of magnitude for much of Earth history (between ∼0.001 and 0.1 bar CO_2_ through the Precambrian (Catling & Zahnle, 2020)) and that RuBisCO catalytic efficiency itself is sensitive to atmospheric CO_2_ /O_2_ levels (Erb & Zarzycki, 2018; Kacar, Hanson-Smith, Adam, & Boekelheide, 2017; Poudel et al., 2020; Riebesell, Revill, Holdsworth, & Volkman, 2000; Schubert & Jahren, 2012; Scott et al., 2007; Tcherkez et al., 2006; Wilkes et al., 2018). Though experimental work has demonstrated the positive correlation between CO_2_ levels and RuBisCO isotopic discrimination (Freeman & Hayes, 1992; Hinga et al., 1994; Schubert & Jahren, 2012; Wilkes et al., 2018), the role of RuBisCO expression, given its CO_2_ sensitivity, has not yet been explored. More broadly, the influences of genetic regulatory processes on ancient biosignatures—otherwise highlighted in geobiological and paleobiological studies of metazoan evolution (Erwin, 2020; Sumner, 2002; Valentine, 1994)—are not often examined in the context of Precambrian microbial metabolisms like carbon fixation. Interpretation of the carbon isotope record should consider the coupled influence of CO_2_ concentration and RuBisCO expression in generating isotopic biosignatures and shaping ancient primary productivity.

To investigate the interplay between RuBisCO expression and CO_2_ concentrations on cyanobacterial physiology and carbon isotopic discrimination, we generated a genetic system to manipulate the expression levels of RuBisCO in the model organism *Synechococcus elongatus* PCC 7942 (hereafter *S. elongatus*).(Bonfil et al., 1998; Gabay, Lieman-Hurwitz, Hassidim, Ronen-Tarazi, & Kaplan, 1998; Maeda, Price, Badger, Enomoto, & Omata, 2000; Omata, Gohta, Takahashi, Harano, & Maeda, 2001; Tchernov et al., 2001). Genetic engineering is facilitated by this strain’s ability to take up and integrate exogenous DNA into its genome by homologous recombination (Taton et al., 2020). *S. elongatus* also possesses the β-carboxysome-based CCM that likely impacted the production of Precambrian carbon biosignatures following the rise of atmospheric oxygen (Hurley et al., 2021; Lyons, Reinhard, & Planavsky, 2014). Thus, *S. elongatus* is a suitable model to study the genetic basis of ancient carbon biosignatures. We engineered its genome with an additional copy of the RuBisCO operon in a chromosomal neutral site to permit genetic manipulation of the RuBisCO operon and confirmed overexpression of RuBisCO under varying CO_2_ levels. Then, to assess whether physiological and subcellular parameters and different CO_2_ concentrations modulate carbon isotopic discrimination in cyanobacteria, we determined growth rate, photosynthetic oxygen evolution rate, and ^13^C/^12^C discrimination of the engineered strain in comparison to wild-type *S. elongatus* under different atmospheric conditions.

## RESULTS

### A second copy of the *rbc* operon results in increased amount of active RuBisCO

The RuBisCO Form IB enzyme in *S. elongatus* is encoded by an operon that includes a CO_2_ -sensitive promoter region (Sengupta et al., 2019) as well as the structural *rbcL* (large subunit) and *rbcS* (small subunit) genes (Vijayan, Jain, & O’Shea, 2011). We designed *S. elongatus* strain Syn02 that harbors the native *rbc* operon and a second copy inserted in the chromosome neutral site 2 (NS2), a site that permits genetic modification without additional indirect phenotypic impact (Andersson et al., 2000; Clerico, Ditty, & Golden, 2007) (**Fig. 1**). Additionally, we generated a control strain (Syn01) whereby RuBisCO is provided solely by an engineered *rbc* operon at its NS2 site (**Table 1**). Attributes of these strains related to carbon fixation, including RuBisCO transcription and protein expression, carboxylase activity, and carbon isotope fractionation, were evaluated under different CO_2_ conditions, including ambient (∼0.04%), 2%, and 5% CO_2_ concentrations. These CO_2_ concentrations were selected to measure the impact of a Precambrian-like atmosphere, which is generally constrained between ∼0.1% to ∼10% CO_2_ for much of the early evolution of cyanobacteria (assuming ∼1-bar total paleopressure) (Catling & Zahnle, 2020).

**Table 1.**
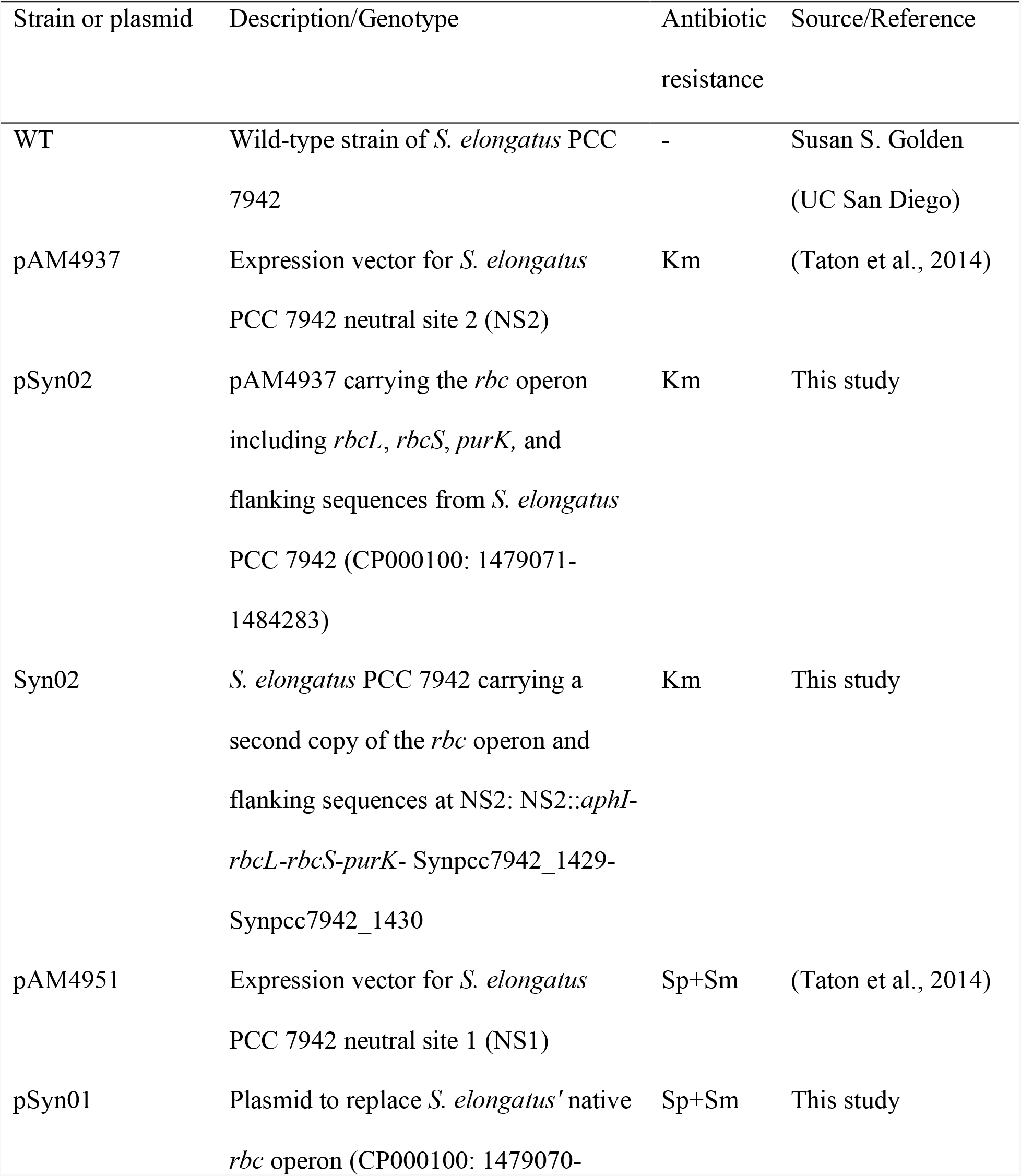
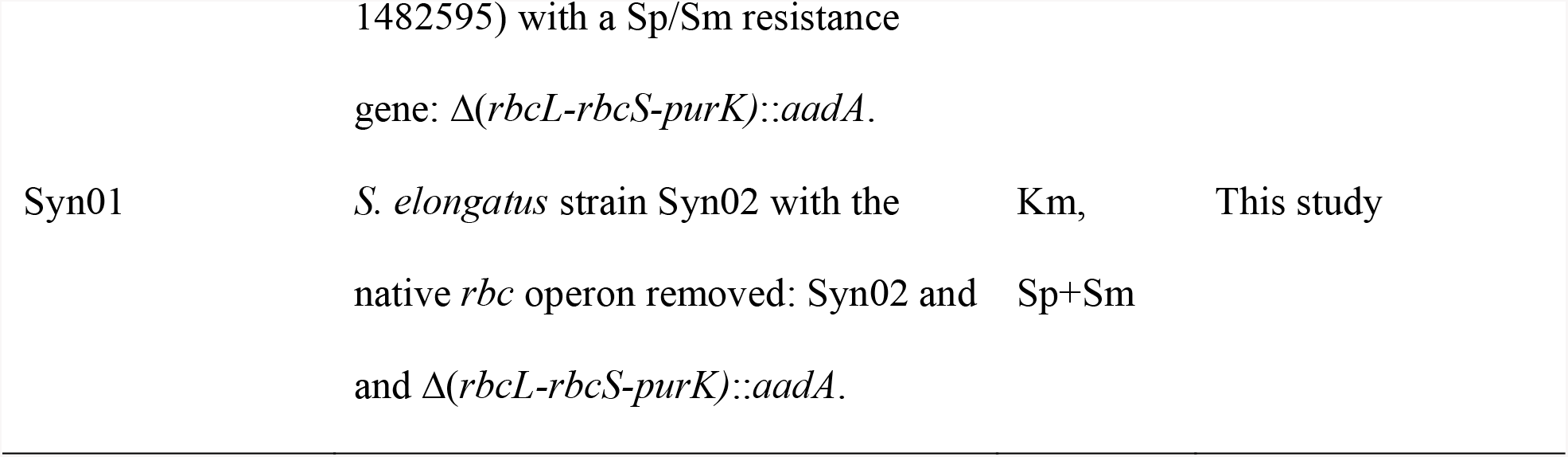
Strains and plasmids used in this study.

**Figure 1.**
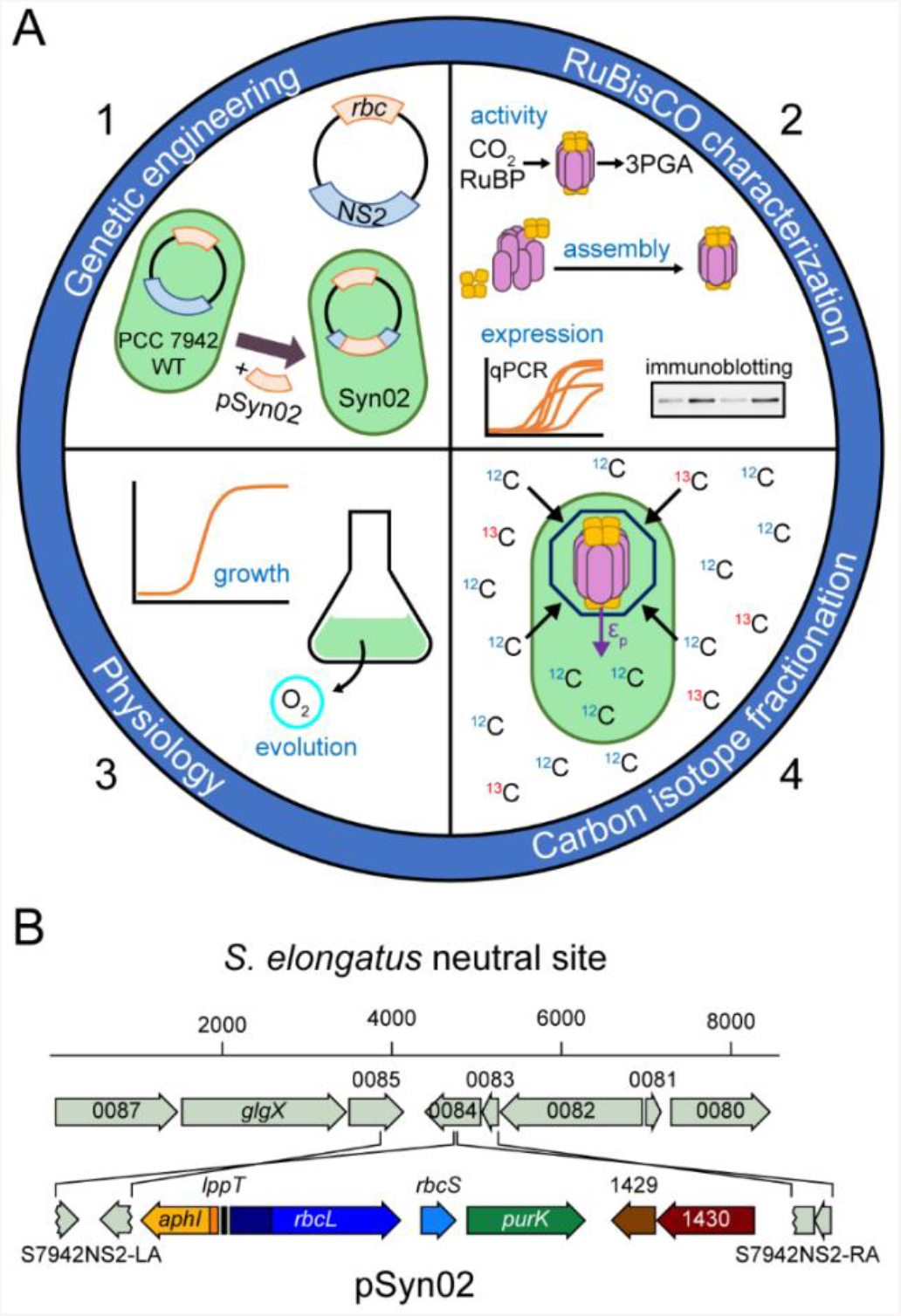
Experimental strategy. **A – Overall methodology.**<u>**1)** Genetic engineering:</u> An additional copy of the *rbc* operon was inserted at the chromosomal NS2 site of *S. elongatu*s PCC 7942 to create the Syn02 strain. <u>**2)** RuBisCO characterization</u> was carried out by investigating (a) catalytic activity, (b) assembly into a hexadecameric complex, and (c) expression at both the transcript and protein levels. <u>**3)** Physiology:</u> *S. elongatu*s WT and Syn02 strains were evaluated by monitoring microbial growth characteristics and rate of photosynthetic oxygen evolution. <u>**4)** Carbon isotope fractionation:</u> The impact of RuBisCO overexpression on biomass δ^13^C and ε_p_ under varying CO_2_ concentrations was assessed. **B** – **Map of the genetic insert in *S. elongates* strain Syn02**. Plasmid pSyn02 was used to insert the *rbc* operon and the *aph1* gene conferring kanamycin resistance in PCC 7942 WT NS2 to generate the Syn02 strain. Crosslines indicate homologous recombination sites and scale bars show DNA fragment sizes (in bp).

We evaluated whether the additional copy of the *rbc* operon in strain Syn02 resulted in increased RuBisCO *rbcL* and *purK* (located downstream of *rbcL* in the operon) transcription under varying CO_2_ levels (ambient, 2%, and 5% CO_2_). Transcription was measured by quantitative reverse-transcription PCR (RT-qPCR), normalized to *secA* and *ppc* reference genes (Hood, Higgins, Flamholz, Nichols, & Savage, 2016; Luo et al., 2019; Szekeres, Sicora, Dragos, & Druga, 2014). We observed that elevated CO_2_ enhanced WT *rbcL* transcription relative by at least ∼5-fold in PCC7942 (*p* < 0.001; **Fig. 2B, Fig. S1B; see Materials and Methods**). *rbcL* transcription was further increased by at least ∼2-fold in Syn02 relative to WT across all tested CO_2_ levels and growth phases, and as high as ∼14-fold in air (*p* < 0.01; **Fig. 2A, Fig. S1A**). Expression of *purK* in Syn02 increased by >2-fold at 2% and 5% CO_2_ relative to WT, but decreased by ∼0.5-fold in air (**Fig. S1C**).

**Figure 2.**
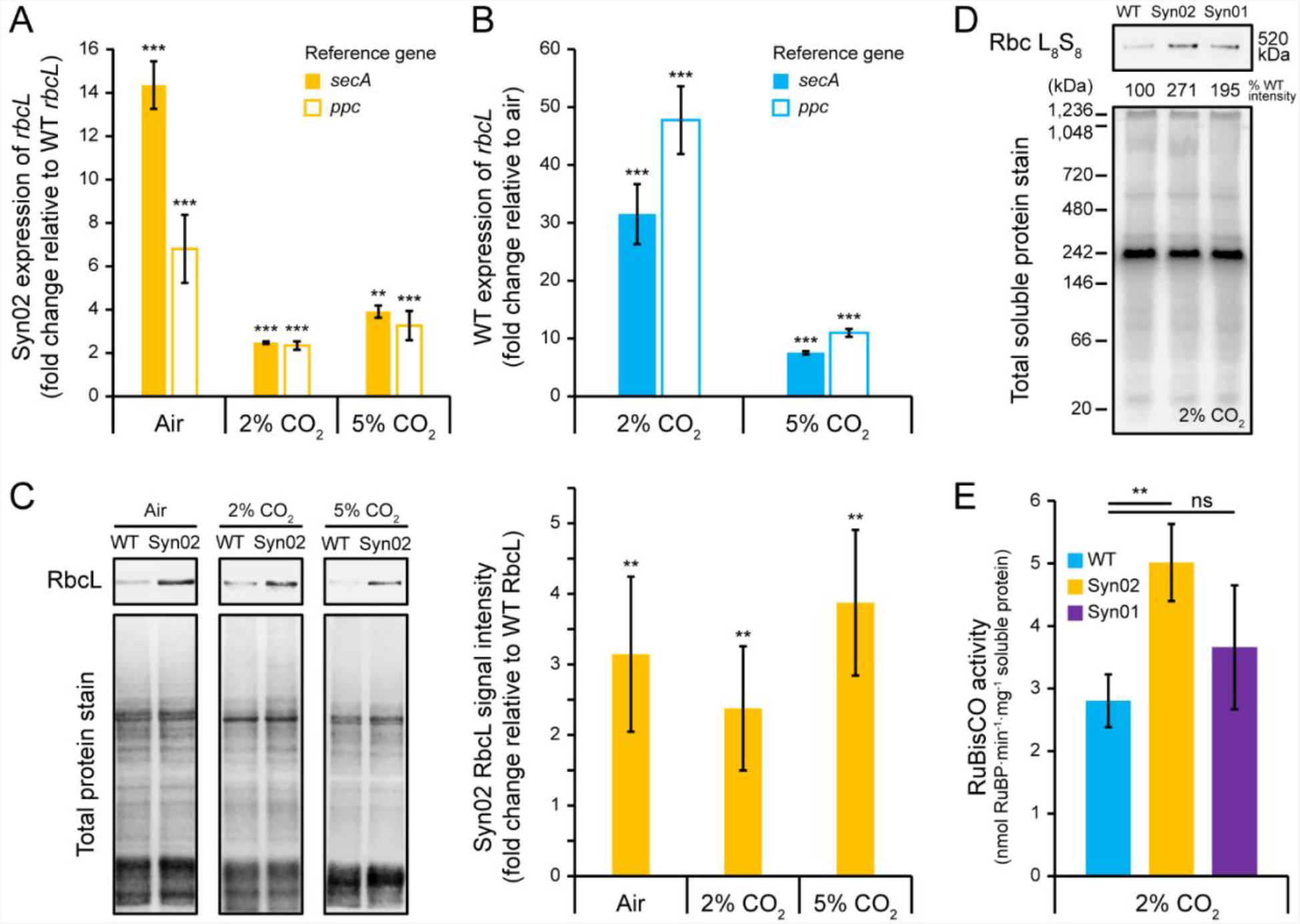
RuBisCO expression, assembly, and activity in *S*. elongatus strains under varying CO_2_ concentrations. **A** – **Expression of Syn02 *rbcL* relative to WT *rbcL***. Asterisks indicate *t*-test results compared to WT *rbcL* expression at the same growth condition. **B – Expression of WT *rbcL* at 2 and 5% CO**_**2**_ **relative to ambient air**. Asterisks indicate *t*-test results compared to WT *rbcL* expression in air. **A-B –** Expression data was measured by RT-qPCR, normalized to reference genes *secA* and *ppc*. Columns represent mean fold expression for three biological replicates. **C – Immunodetection of RbcL**. (*left*) Western blot showing RbcL protein detected by anti-RuBisCO antibody and total protein stain from crude cell lysates. (*right*) RbcL percentage signal intensities were normalized to that for the total soluble protein load. Columns represent mean RbcL intensities for six biological replicates, relative to mean values for WT cultures at the same growth condition. Asterisks indicate *t*-test results compared to WT RbcL. **D – Immunodetection of assembled RuBisCO**. Western blot showing the proper assembly of RbcL and RbcS into the L_8_ S_8_ hexadecameric complex L_8_ S_8_ (520 kDa), detected by anti-RbcL antibody. Rbc L_8_ S_8_ percentage signal intensity was normalized to that for the total soluble protein load, and is shown relative to WT. **E – Lysate RuBisCO activity**. Activity was measured by the RuBP consumption rate, normalized to total soluble protein content of cell lysate. Columns represent the mean activity of three biological replicates. **A-C, E –** Asterisks indicate *t*-test results for pairwise comparison indicated by horizontal line. Error bars on all graphs indicate 1 SD. ns – not significant; ** – *p* <0.01; *** – *p* <0.001.

To determine *rbcL* overexpression at the protein-level, we quantified RbcL protein from crude cell lysates by Western blot using rabbit anti-RbcL antibody (see **Materials and Methods**). In agreement with RuBisCO overexpression indicated by the RT-qPCR results, densitometric analyses revealed a mean ∼2 to 4-fold increase of RbcL protein in Syn02 relative to WT across all tested CO_2_ concentrations (*p* < 0.01; **Fig. 2C**). Finally, we confirmed proper assembly of the large (L) and small (S) subunits into the RuBisCO L_8_ S_8_ complex in Syn02 as well as the Syn01 control strain by native protein electrophoresis and detection by anti-RbcL antibody. At 2% CO_2_, we found a ∼3-fold increase in assembled RuBisCO protein for Syn02 and ∼2-fold increase for Syn01 relative to WT for cultures grown in 2% CO_2_ (**Fig. 2D**).

Finally, we tested whether RuBisCO overexpression in Syn02 resulted in an increased amount of active RuBisCO by determining the total carboxylase activity of cell lysates. Because we detected the smallest increase in transcript levels for Syn02 relative to WT under 2% CO_2_, we selected this growth condition to provide a reasonable lower bound on Syn02 RuBisCO activity. We measured a mean ∼1.8-fold (*p* < 0.01) increase in Syn02 lysate RuBisCO activity relative to WT (**Fig. 2E**). Despite the modest increase in assembled RuBisCO detected for the control strain, Syn01, no significant difference in activity was found between Syn01 and WT cultures grown in these same conditions.

### RuBisCO overexpression does not strongly influence growth rate or photosynthetic activity

Following confirmation that Syn02 overexpresses active RuBisCO, we compared growth rates of WT and Syn02 *S. elongatus* strains at different CO_2_ levels to identify potential downstream physiological effects of increased RuBisCO protein. WT and Syn02 both exhibited ∼2.5-fold faster growth rates under 2% and 5% CO_2_ compared to ambient air (*p* < 0.001; **Fig. 3; Table 2**). Syn02 exhibited a slight ∼1.1-fold increase in growth rate in ambient air relative to WT (*p* < 0.05). No difference in growth rate was observed between the two strains under 2% and 5% CO_2_. The carrying capacity (maximum cell density measured at OD_750_ across the total growth period) for cultures varied between different atmospheric conditions, with a maximum carrying capacity of OD_750_ ≈ 8.4 reached under 2% CO_2_. No significant difference in carrying capacity was found between WT and Syn02 cultures grown under the same atmospheric conditions. These experiments were replicated with unsparged cultures, yielding consistent results (**Table S1**).

**Table 2.** Growth parameters and oxygen evolution rates of *S. elongatus* strains under varying CO_2_ concentrations^a^.

| Strain | Atmosphere | Growth rate | Carrying capacity (OD<sub>750</sub>) | Oxygen evolution rate |
| --- | --- | --- | --- | --- |
|  |  | (h<sup>-1</sup>) |  | (nmol O<sub>2</sub>·h<sup>-1</sup>·μg<sup>-1</sup> chlorophyll *a*) |
| WT | Air | 0.017 ± 0.001 | 6.8 ± 0.7 | 340 ± 30 |
|  | 2% CO<sub>2</sub> | 0.043 ± 0.002 | 8.4 ± 0.5 | - |
|  | 5% CO<sub>2</sub> | 0.047 ± 0.005 | 5.4 ± 0.3 | - |
| Syn02 | Air | 0.020 ± 0.001* | 7.6 ± 0.5 | 360 ± 30 |
|  | 2% CO<sub>2</sub> | 0.046 ± 0.001 | 8.33 ± 0. | - |
|  | 5% CO<sub>2</sub> | 0.040 ± 0.001 | 5. ± 0.4 | - |

<sup>a</sup>Values are means of three biological replicates ± 1 SD. Asterisks indicate *t*-test result from comparison with WT for the same atmospheric condition, \* – *p* < 0.05.

**Figure 3.**
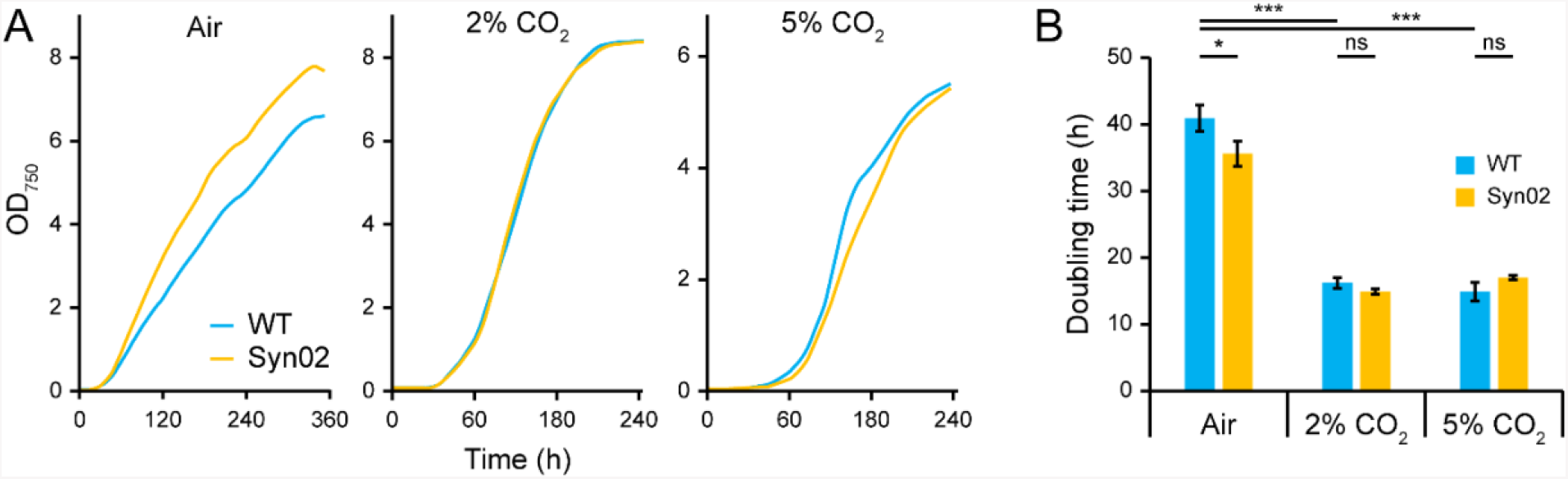
Growth of *S. elongatus* strains. A – Growth profiles. Cultures of each strain were maintained in ambient air or under 2% or 5% CO_2_. Smoothed profiles were generated from mean optical density values (measured at 750 nm, OD_750_)for three replicate cultures per growth condition. **B – Doubling times**. Columns represent the mean doubling time of three replicates. Error bars on all graphs indicate 1 SD, and asterisks indicate *t*-test results for pairwise comparisons indicated by horizontal lines. ns – not significant; * – *p* <0.05; *** – *p* < 0.001.

To further test for differences in photosynthetic activities between the WT and Syn02 strains, we measured their oxygen evolution rates in ambient air. After brief incubation in the dark, culture samples were exposed to saturated light in an oxygen electrode chamber to detect levels of molecular oxygen. Oxygen evolution rates were normalized to chlorophyll *a* concentrations, following Zavřel et al. (2015). We did not observe a significant difference in mean oxygen evolution rate between WT and Syn02 (340 ± 30 O_2_ ·h^-1^·µg^-1^ chlorophyll *a* and 360 ± 30 O_2_ ·h^-1^·µg^-^ _1_ chlorophyll *a*, respectively) (**Table 2**).

### *S. elongatus* ^13^C/^12^C fractionation is insensitive to RuBisCO overexpression

We tested the combined influence of RuBisCO overexpression and CO2 concentration on the magnitude of whole-cell ^13^C/^12^C isotopic discrimination in *S. elongatus*. The isotopic discrimination associated with photosynthetic CO_2_ fixation (ε_p_) was calculated from measured δ^13^C_biomass_ and headspace δ^13^C_CO2_ in our experimental system, as described in **Materials and Methods**. We observed that increased CO_2_ concentration (2% CO_2_) increases the degree of *S. elongatus* ^13^C/^12^C discrimination relative to air. Mean ε_p_ values for WT and Syn02 strains grown in ambient air were 4.8 ± 0.3‰ and 5.1 ± 0.4‰, respectively (**Fig. 4; see Supplementary Information, Table S2** and **S3** for δ^13^C and pH measurement valuesvalues). At 2% CO_2_, mean ε_p_ values for WT and Syn02 strains were 23.75 ± 0.07‰ and 23.7 ± 0.5‰, respectively. The difference between WT and Syn02 ε_p_ values at both tested CO_2_ conditions were found to be insignificant (*p* > 0.05). These results demonstrate that the increased RuBisCO transcriptional levels and total carboxylasecarboxylase activity measured for Syn02 in our experiments does not significantly impact whole-cell *S. elongatus* ^13^C/^12^C discrimination, both at ambient and elevated CO_2_ concentrations.

**Figure 4.**
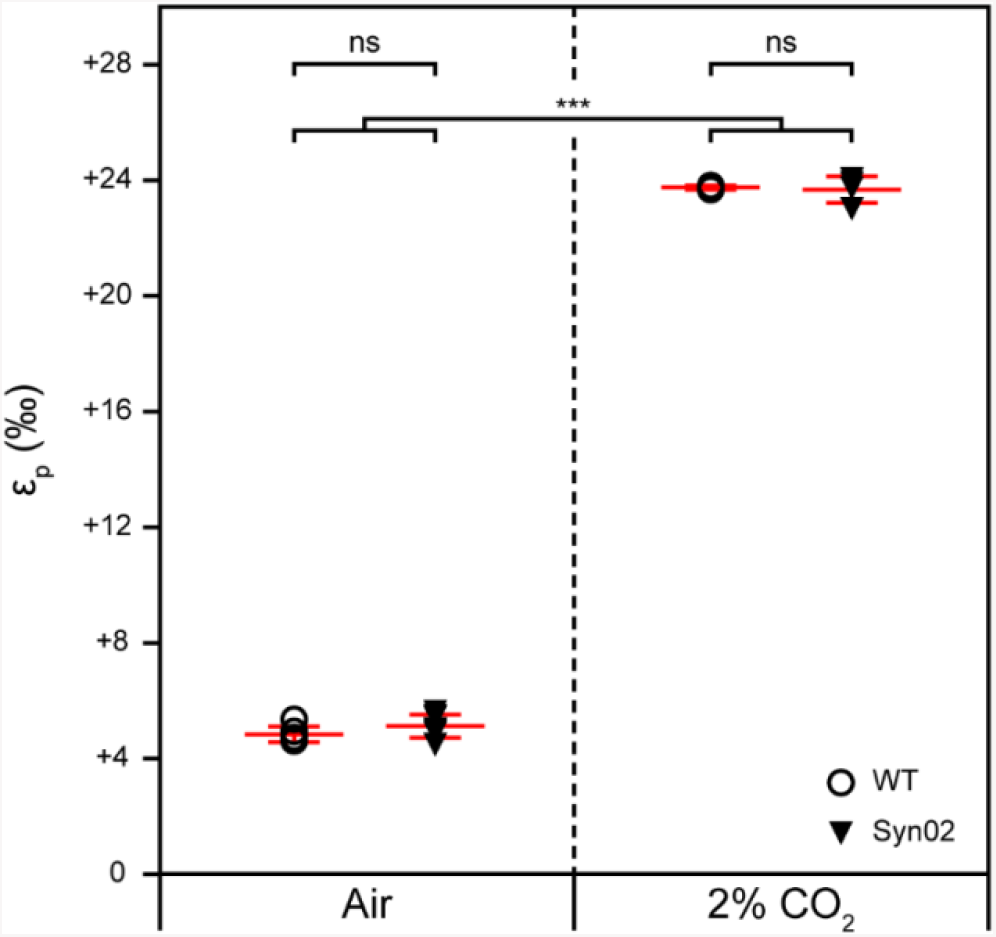
^13^C/^12^C discrimination associated with photosynthetic CO_2_ fixation (ε_p)_ of *S. elongatus* strains under ambient and 2% CO_2_ concentrations. ε_p_ values are calculated relative to measured δ^13^C_CO2_ (Table S3; see **Materials and Methods** for calculation). Mean (long horizontal bars) and 1 SD (vertical error bars) are shown in red, calculated for seven (air) and four (2% CO_2_) biological replicates. Asterisks indicate *t*-test results for pairwise comparisons indicated by horizontal lines. ns – not significant; *** – *p* < 0.001.

## DISCUSSION

Evolutionary biologists have long debated the role of genetic regulation versus protein-level variation in evolutionary selection (Carroll, 2005; Fay & Wittkopp, 2008; Olson-Manning, Wagner, & Mitchell-Olds, 2012; Taylor, Shepherd, Jackson, & Silby, 2022; Wilson, Maxson, & Sarich, 1974). How such molecular and genetic variation leads to evolutionary change is often overlooked in geobiological evaluation of ancient microbial biosignatures due to the challenges involved with disentangling their effects from other variables. Laboratory experimental systems, though admittedly incapable of capturing the full complexity of ancient, natural systems, are necessary to understand the genetic factors underpinning the oldest records of life. In this study, we developed an engineered laboratory system to investigate the coupled impact of elevated CO_2_ levels and RuBisCO expression on cyanobacterial physiology and carbon isotope discrimination. We tested whether the overexpression of RuBisCO influences *S. elongatus* growth, photosynthetic oxygen evolution rate, and carbon isotopic signatures at different CO_2_ atmosphere concentrations by generating an engineered *S. elongatus* strain Syn02 that harbors a second copy of the *rbc* operon at NS2.

We confirmed overexpression of RuBisCO in Syn02 by combining transcriptional analyses, protein quantification, and assays of total protein carboxylation activity. Our results show elevated RuBisCO transcription in *S. elongatus* strain Syn02 relative to WT at all tested CO_2_ concentrations (ambient air, 2%, and 5% CO_2_) (**Fig. 2A**), with a ∼2-fold increase in active, assembled RuBisCO protein measured under 2% CO_2_ (**Fig. 2D and 2E**). We also found that higher levels of CO_2_ result in increased RuBisCO transcription for both WT and Syn02 strains relative to ambient air, with the greatest fold change increase detected at 2% CO_2_. This result is in agreement with a previous study reporting a regulatory element in the *S. elongatus* PCC 7942 *rbcL* operon that regulates the expression of RbcL with increasing CO_2_ (Sengupta et al., 2019).

Our experimental approach also enabled us to detect potential upper bounds on the ability of *S. elongatus* to overexpress RuBisCO under a variety of CO_2_ conditions. First, increased *rbcL* transcription in Syn02 relative to WT did not always result in a proportional increase in translated RuBisCO protein. For example, even though Syn02 showed a ∼14-fold increase in *rbcL* transcript relative to WT in air (**Fig. 2A**), total Syn02 RbcL protein only increased by ∼3-fold under these same conditions (**Fig. 2C**). These results indicate that other physiological factors, possibly including translation efficiency or expression of ancillary proteins in our experimental system, may ultimately limit RuBisCO expression beyond transcription. Further, it is important to consider whether all translated RuBisCO protein is catalytically active. Previous studies have shown that overexpression of RuBisCO in cyanobacteria generally produces an increased pool of active carboxylase (Atsumi, Higashide, & Liao, 2009; Iwaki et al., 2006; Lechno-Yossef et al., 2020; Liang & Lindblad, 2017). Indeed, for cells grown in 2% CO_2_, we found that nearly all overexpressed RuBisCO protein was likely to be assembled and catalytically active (**Fig. 2C** and **2E**). Future work may determine whether this proportion of active, overexpressed RuBisCO holds for cells cultivated at other CO_2_ conditions. Finally, we observed that the transcriptional impact of a second *rbc* copy in Syn02 was dampened at elevated CO_2_ relative to air (**Fig. 2A**), suggesting a potential limit to the combined expression-level effects of increased *rbc* copy number and increased CO_2_. Altogether, our findings suggest the presence of physiological constraints that may be limiting RuBisCO overexpression at both transcript and protein levels, the exact mechanism of which awaits further characterization. Similar approaches investigating other model microbial systems would clarify the degree to which these observations might extend to cyanobacteria gene regulation in the past.

We investigated whether the overexpression of RuBisCO in strain Syn02 impacted the growth characteristics of *S. elongatus* and, by that manner, might have the potential to influence isotopic discrimination (Bidigare et al., 1997; Laws et al., 1997; Wilkes et al., 2018). Though various culture conditions including pH, temperature, CO_2_ concentration, nutrient availability, and light intensity may substantially affect the growth characteristics of cyanobacteria (Kuan, Duff, Posarac, & Bi, 2015; Rillema, MacCready, & Vecchiarelli, 2020; Ungerer et al., 2018; Yu et al., 2015), the influence of genetic factors, including the regulation of RuBisCO expression, is less well known. Previous studies targeting the correlation between RuBisCO overexpression and cyanobacterial physiology have yielded mixed results. For instance, faster growth rates and oxygen evolution rates were observed for an engineered *Synechocystis* PCC 6803 strain that overexpresses RuBisCO (Liang & Lindblad, 2017), as well as in *S. elongatus* upon co-overexpression of its phosphoribulokinase (Kanno, Carroll, & Atsumi, 2017). However, in *Synechococcus* sp. PCC 7002, the overexpression of RuBisCO did not alter growth rate (De Porcellinis et al., 2018). The impact of RuBisCO upregulation on cyanobacterial growth appears to be species/strain-specific. Our data shows that RuBisCO overexpression in *S. elongatus* PCC 7942 strain Syn02 results in a minor increase in growth rate in ambient air relative to WT, and no significant difference in growth or oxygen evolution rates was observed at elevated CO_2_ levels. Our results indicate that further study is needed to fully understand the coupled interaction of different environmental conditions and genetic backgrounds in determining cyanobacterial growth parameters of relevance to carbon biosignature generation.

We found that the degree of RuBisCO overexpression generated in Syn02 did not measurably impact *S. elongatus* carbon isotope discrimination, both in ambient air and at elevated CO_2_ (**Fig. 4**). Rather, the ε_p_ variability observed in our experiments stemmed solely from variable CO_2_, an effect that has previously been observed empirically for a variety of autotrophs (Freeman & Hayes, 1992; Hinga et al., 1994; Schubert & Jahren, 2012; Wilkes et al., 2018), including for cyanobacteria in particular (Eichner et al., 2015; Hurley et al., 2021). Other studies have shown that photoautotrophs grown under a lower pH, and thus a higher proportion of CO_2_, also exhibit higher carbon fractionation (Mizutani & Wada, 1982; Roeske & O’Leary, 1984; Wang, Yeager, & Lu, 2016; Yoshioka, 1997). In our experiments, ε_p_ values were increased by ∼19‰ for WT and Syn02 strains in 2% CO_2_ relative to air, the latter of which was carbon limiting in our experimental system (see **Supplementary Information**). Theoretically, under high CO_2_ concentrations, RuBisCO carbon fixation is presumed to be rate-limiting and its intrinsic kinetic isotope effect is expressed, maximizing ε_p_ (Bidigare et al., 1997; Hayes, 1993; Schubert & Jahren, 2012; Wilkes et al., 2018). Under high CO_2_ availability, CCM modules have been found to be downregulated in cyanobacteria, including *S. elongatus* PCC 7942 (Burnap, Hagemann, & Kaplan, 2015; Schwarz et al., 2011; Woodger, Badger, & Price, 2003). Indeed, the mean ε_p_ ≈ 24‰ value we observed under high inorganic carbon availability is within the ^13^C/^12^C isotopic effect range previously measured for cyanobacterial Form IA and IB RuBisCO at saturating CO_2_ (∼22–24‰) (Guy et al., 1993; Scott et al., 2007). Nevertheless, we cannot rule out the impact of upstream cellular modules in significantly shaping the whole-cell, isotopic signatures observed in our experimental system, which recent work has demonstrated can exert important isotopic effects under a variety of growth conditions (Burnap et al., 2015; Eichner et al., 2015; Hurley et al., 2021; Wilkes & Pearson, 2019). Though we did not quantify the isotopic contributions of these other intracellular factors in our engineered system, it seems apparent that the increased RuBisCO expression and total protein carboxylase activity in Syn02 neither changed the ^13^C/^12^C substrate pool in the enzyme’s vicinity nor measurably altered the potential impact of other cellular modules on carbon isotope discrimination, even at ambient CO_2_ levels.

The range of ε_p_ values measured for both *S. elongatus* WT and Syn02 strains under elevated CO_2_ levels plausibly experienced by ancient cyanobacteria falls within the ∼8 to 24‰ ε_p_ range inferred from the Precambrian carbon isotope record following the rise of atmospheric oxygen (Hurley et al., 2021; Krissansen-Totton et al., 2015). Our data suggest that potential overexpression of RuBisCO by past cyanobacterial primary producers likely does not help explain the distribution of Precambrian carbon biosignatures. Rather, it is more probable that other environmental and intracellular factors, including but not limited to CO_2_ levels, growth rates, temperature, and CCM effects, are more significant determinants of the isotopic compositions of organic carbon. We note in particular that less well-characterized RuBisCO forms (i.e., Forms IC, ID, and II) that are found in proteobacteria and marine phytoplankton can exhibit significantly smaller fractionation factors than that observed for forms associated with land plants and cyanobacteria, and perhaps were more dominant in ancient ecosystems (Boller et al., 2011; Robinson et al., 2003; Thomas et al., 2019). The full potential breadth of geochemical carbon fixation signatures requires further study (Garcia et al., 2021), which can be significantly augmented by the integration of taxonomically undercharacterized systems, the regulatory dynamics of the carbon fixation protein network beyond RuBisCO, and laboratory reconstructions of ancient carbon-fixing enzymes (Kedzior et al., 2022). Our results underscore that the integration of synthetic biology, metabolic engineering, and geochemistry can offer new insights into the study and interpretation of biogeochemical reservoirs at the global scale. Integrative laboratory approaches are crucial to establish the extent to which the modern biological carbon isotope fractionation properties can be correlated with evidence for microbial activity in the Earth’s deep past.

## MATERIALS AND METHODS

### Cyanobacterial growth and maintenance

*S. elongatus* PCC 7942 strains were cultured in BG-11 medium (Rippka, Deruelles, Waterbury, Herdman, & Stanier, 1979) as liquid cultures or on agar plates (1.5% (w/v) agar and 1 mM Na_2_S_2_O_3_·5H_2_O). For recombinant strains, liquid and solid media were supplemented with appropriate antibiotics: 2 μg·ml^-1^ Spectinomycin (Sp) plus 2 μg·ml^-1^ Streptomycin (Sm), 5 μg·ml^-1^ Kanamycin (Km). The cyanobacterial growth was measured by optical density at 750 nm (OD_750_).

The strains were archived at -80 °C in 15% (v/v) glycerol. The 2-ml glycerol stocks were rapidly thawed and inoculated in liquid BG-11 (supplemented with antibiotics as needed). Liquid cultures were maintained in a Percival environment chamber (Cat. No. I36LLVLC8) fitted to a CO_2_ gas cylinder input, permitting monitoring and control of internal atmospheric CO_2_. Cultures were shaken at 120 rpm at 30 °C under continuous low illumination of 45 µmol photon·m^-2^·s^-1^, and ambient air, until they reached an OD_750_ between 0.4 and 0.6. These cultures were then used to inoculate fresh cultures that were grown using similar conditions but under moderate illumination of 80 µmol photon·m^-2^·s^-1^ (following (Smith & Williams, 2006)) and sparged at selected CO_2_ concentrations (ambient air, 2%, or 5% CO_2_ ; cultures were also grown without sparging for carbon isotope fractionation experiments and to assess differences in growth rate; **Table S1**). Cultures were sampled at an OD_750_ of ∼7 to 7.5 for most subsequent experiments, as described below. Cultures were sampled at an earlier OD_750_ of 0.5-1.5 for carbon isotope fractionation measurement and for additional RT-qPCR experiments (**Fig. S1**).

### Genetic engineering of cyanobacteria

A recombinant strain of *S. elongatus* was constructed by natural transformation using standard protocols (Clerico et al., 2007) and the plasmids and methods described below (**Table 1**). To construct the plasmid pSyn02, pAM4937 was digested with *Swa*I to release the *ccdB* toxic gene and produce a plasmid backbone that contains the pBR322 *E. coli* origin of replication, the base of mobilization site for conjugal transfer, the *aph1* gene conferring kanamycin resistance, and sequences for homologous recombination into *S. elongatus* chromosome at NS2. The *rbc* operon was amplified from *S. elongatus* PCC 7942 gDNA with primers F01 and R01 (**Table S4**) containing 20-nucleotide sequences that overlap with pAM4937 backbone. The resulting DNA fragments were assembled using the GeneArt™ Seamless Cloning and Assembly Kit (Invitrogen, Cat. No. A13288). pSyn02 was used to insert the *rbc* operon into NS2 of the wild-type genome in the strain PCC 7942 through homologous recombination to create the strain of *S. elongatus*, Syn02, carrying two copies of the *rbc* operon. Plasmid pSyn-01 was constructed by using two primer pairs F02/R02 and F03/R03 (**Table S4**), 1) to amplify a fragment of pAM4951 that contains the *E. coli* origin of replication and the site for conjugal transfer, and 2) to amplify the *aadA* gene conferring spectinomycin/streptomycin resistance. Native *rbc* operon flanking sequences were amplified from the *S. elongatus* PCC 7942 gDNA with the primer pairs F04/R04 and F05/R05 (**Table S4**) containing 20-nucleotide sequences that overlap with the pAM4951 fragments.

Transformation was carried out after growing the WT strain in liquid culture at 30 °C with shaking (120 rpm) and a light intensity of 80 µmol photon·m^-2^·s^-1^ until an OD_750_ ∼0.7. The cells were prepared for transformation according to the protocol by Clerico et al. (2007) and plated on BG-11 agar containing the appropriate antibiotic(s) for recovery. Subsequently, the colonies were picked using sterile pipette tips, patched onto BG-11 agar containing the appropriate antibiotic(s), and further incubated to ensure complete chromosome segregation (i.e., incorporation of the *trans*-gene into all chromosomes). The patched transformants were screened using colony PCR using the primers F06/R06 and F07/R07 and the genotypes of the engineered strains were confirmed by Sanger sequencing using the primers F08/R08 **(Fig. S3, Table S4**)

### Extraction of total RNA and proteins

Cultures were collected during the exponential growth phase. Cells were pelleted by centrifugation at 4,700 × g for 10 min at room temperature and resuspended in 10 mL of TE buffer (10 mM Tris, pH 8.0, 1 mM EDTA). To prepare the crude cell lysate, 8 mL of the cell suspension were pelleted by centrifugation at 4,700 × g for 10 min at room temperature, resuspended in 500 µL of hot (pre-warmed to 95 °C) TE buffer supplemented with 1% (w/v) SDS and incubated at 95 °C for 10 min. The mixture was sonicated at 40% amplitude for 3 × 10 sec with 10 sec intervals and the cell debris was centrifuged at 17,000 × g for 10 min at room temperature. The supernatant was collected and stored at -80 °C. Total RNA was extracted from the remaining 2 mL of the cell suspension using the RNeasy® Protect Bacteria Mini Kit (QIAGEN, Cat. No. 74524) following the manufacturer instructions.

### Analysis of *rbc* operon expression by RT-qPCR

Total RNA was quantified with a NanoDrop spectrophotometer and 1 µg of RNA was treated with the amplification grade deoxyribonuclease I (Invitrogen, Cat. No. 18068-015). 10 µL of DNase I-treated RNA was then used for reverse transcription (RT) performed with the SuperScript™ IV First-Strand Synthesis System (Invitrogen, Cat. No. 18091050). The four pairs of qPCR primers (listed in **Table S4**) were designed with Primer3Plus (http://www.bioinformatics.nl/cgi-bin/primer3plus/primer3plus.cgi). Both reference genes have previously been shown to be stably expressed under diverse conditions in *S. elongatus* (Luo et al., 2019). The quality of cDNA and primer specificity was assessed by PCR using cDNA templates (RT positive reactions), RT negative controls, and the qPCR primers. The analysis of gene expression levels was performed in a real-time thermal cycler qTOWER^3^ G (Analytik Jena AG), equipped with qPCRsoft software, using the cycles: 50 °C/2 min, 95 °C/2 min, 40 × (95 °C/15 sec, 60 °C/1 min). The relative expression of the *rbc* operon genes (*rbcL* and *purK*) was calculated as the average fold change normalized to *secA or ppc* reference genes using the delta-delta Ct method. The experiment was carried out using three biological replicates and three technical replicates.

### Analysis of RbcL protein by Western blot

Total protein concentration in the crude cell lysates was measured using the Pierce™ BCA Protein Assay Kit (Thermo Scientific, Cat. No. 23225). The lysates were loaded in the amount of 5 µg of total protein in Laemmli sample buffer onto a 6% (v/v) polyacrylamide stacking gel. Proteins were electrophoresed in a 12% polyacrylamide resolving gel in TGS buffer and blotted in transfer buffer onto a PVDF membrane. Total protein load in each sample was visualized by Revert™ 700 Total Protein Stain (LI-COR Biosciences, Cat. No. 926-11011) and used for RbcL signal normalization. Detection of RbcL was performed by overnight incubation of the membrane at 4 °C with rabbit anti-RbcL antibody (Agrisera, Cat. No. AS03 037), 1:5000 in TBST with 5% non-fat milk, followed by one-hour incubation at room temperature with IRDye® 800CW goat anti-rabbit IgG secondary antibody (LI-COR Biosciences, Cat. No. 926-32211), 1:20,000 in Intercept® (TBS) blocking buffer (LI-COR Biosciences, Cat. No. 927-60001) with 0.1% (v/v) Tween-20 and 0.01% (w/v) SDS. Both the total protein load and the amount of RbcL in each sample were documented with Odyssey® Fc Imaging System (LI-COR Biosciences, Cat. No. 2800-03) at the near-infrared detection mode. The images were acquired using Image Studio™ software. The densitometric analysis of RbcL signal intensity, normalized to total protein load, was performed with Quantity One® software (Bio-Rad) for six biological replicates. The amount of RbcL produced by each replicate of the Syn02 strain culture was compared to the averaged level of RbcL in the WT PCC 7942 replicate cultures and expressed as the averaged percent of RbcL synthesized by the WT strain.

### Assembly of RuBisCO subunits

Assembly of the RuBisCO large and small subunits into a hexadecameric complex in each strain was evaluated by native gel electrophoresis and immunodetection. Samples were collected during the exponential growh phase. Cells grown under 2% CO_2_ were pelleted by centrifugation at 4,700 × g for 10 min at room temperature and resuspended in 400 µL of native lysis buffer (50 mM Tris, pH 8.0, 150 mM NaCl, 1 mM EDTA, 10% (v/v) glycerol) supplemented with 5 mM DTT, 100 µg/mL lysozyme from chicken egg white, and 1% (v/v) Halt™ Protease Inhibitor Cocktail (Thermo Scientific, Cat. No. 78430). The cell suspensions were incubated at 30 °C for 15 min and subjected to five consecutive freeze-thaw cycles (10 min at -80 °C followed by 5 min at 30 °C), then were sonicated on ice for 3 minutes at 30% amplitude (2-sec on/off intervals), centrifuged at 17,000 × g for 15 min at 4 °C. The concentration of total soluble proteins in the lysates was determined with a Pierce™ BCA Protein Assay Kit. The lysates were adjusted to 5 µg of total soluble proteins in native sample buffer and then loaded onto a 4-20% Mini-PROTEAN® TGX™ Precast Protein Gel (Bio-Rad, Cat. No. 4561094). Protein electrophoreses were performed in TG buffer (60 mM Tris, 192 mM glycine) at 100 V for 4 h at 4 °C and blotted in transfer buffer (48 mM Tris, pH 9.2, 39 mM glycine, 0.04% (w/v) SDS) onto a nitrocellulose membrane. After three 10-min washes in wash buffer (48 mM Tris, pH 9.2, 39 mM glycine, 20% (v/v) methanol), total protein load in each sample was visualized by Revert™ 700 Total Protein Stain and used for the normalization of RuBisCO complex quantity. Immunodetection of the RuBisCO complex was performed with the same primary and secondary antibodies that were used to analyze the level of RbcL, as described above.

### Catalytic activity of RuBisCO

The activity of RuBisCO in cyanobacterial lysates was measured using a spectrophotometric coupled-enzyme assay that links this activity with the rate of NADH oxidation (Kubien, Brown, & Kane, 2011). The cyanobacterial strains were cultured under 2% CO_2_, collected, and pelleted as described above. The pellets were resuspended in 1 mL of ice-cold lysis buffer (50 mM EPPS, 1 mM EDTA, 2 mM DTT, pH 8.0) and transferred into 2 mL screw-capped tubes with Lysing Matrix B (MP Biomedical) for lysis by bead beating using FastPrep-24™ 5G bead beater (MP Biomedical) with 4 m/sec for 10 sec, followed by 2-min incubation on ice, repeated six times. The cell lysates were transferred to new Eppendorf tubes to remove the beads and unbroken cells and to pellet the thylakoid membrane by centrifugation at 10,000 × g for 1 min and at 20,000 × g for 30 min at 4 °C, sequentially. The resulting clear supernatants containing cytosolic soluble proteins, including phycobiliproteins and RuBisCO, were used to determine protein concentration by Pierce™ BCA Protein Assay Kit and to measure RuBisCO activity by employing an assay adapted from Kubien et al. (2011). The assay buffer (100 mM HEPES, 25 mM MgCl_2_, 1 mM EDTA, pH 7.6) was used considering the high Michaelis constant for CO_2_ (K_C_) for cyanobacterial RuBisCO. 20 µL of cell lysates were preincubated in the assay mix (with 5 mM NaHCO_3_) at 25 °C for activation before initiating the reaction by adding synthesized ribulose 1,5-bisphosphate (RuBP) according to Kane et al. (1998). The absorbance at 340 nm was monitored using a Synergy H1 plate reader (BioTek). RuBisCO activity was reported as RuBP consumption rate normalized to total soluble protein content. The assay was performed for three biological replicates.

### Cyanobacterial growth measurements

OD_750_ values were plotted as a function of time and analyzed in R with the Growthcurver package. Growth curve data was fitted to the standard form of the logistic equation to calculate growth parameters including growth rate, doubling time, and carrying capacity (Sprouffske & Wagner, 2016). Each strain was grown in triplicate for every condition.

### Photosynthetic oxygen evolution rate

*S. elongatus* strain photosynthetic activity was assayed using a Clark-type oxygen electrode chamber to measure the level of molecular oxygen produced in cyanobacterial cultures. Cells were grown in 50 mL of BG-11 at 30 °C, illumination of 80 µmol photon·m^-2^·s^-1^, shaking at 120 rpm, in ambient air, and with culture sparging. The samples were collected from triplicate cultures during the exponential growth phase, pelleted by centrifugation at 4,700 × g for 10 min at room temperature, and resuspended in fresh BG-11 to an OD_750_ of ∼1.0. Oxygen evolution rates were normalized to chlorophyll *a* concentration (following (Liang & Lindblad, 2017) using the protocol by Zavřel et al. (2015). The remaining suspension was incubated in the dark for 20 min with gentle agitation. Samples from each suspension, prepared in three technical replicates, were analyzed in an oxygen electrode chamber under saturated light, using the Oxygraph+ System (Hansatech Instruments) equipped with the OxyTrace+ software. Oxygen evolution rate was monitored for 10 min and expressed as nanomoles of molecular oxygen evolved per hour per microgram of chlorophyll *a*.

### Carbon isotope fractionation in bulk cyanobacterial biomass

Carbon isotope fractionation experiments were conducted under ambient and 2% CO_2_ conditions. Cyanobacteria were cultured in shaken flasks (all other conditions as described above) to an OD_750_ of 0.5 to 1.5 (seven and four biological replicates per strain for experiments conducted in air and 2% CO_2_, respectively). Cells were pelleted by centrifugation at 4,700 × g for 10 min at room temperature and washed in 10 mL of 10 mM NaCl. The bacteria were resuspended in 1 mL of 10 mM NaCl and transferred to Eppendorf tubes. After centrifugation at 4,700 × g for 10 min at room temperature, the supernatants were completely removed, the pellets were dried in opened tubes in a laboratory oven at 50 °C for 2 days, and the resultant dried biomass samples were transferred into tin capsules. The pH of growth medium aliquots was measured at the beginning (sampled prior to inoculation) and end of cyanobacterial culture (sampled from the supernatant of pelleted cells, see above). Aliquots sampled at the end of the culture period were then filter-sterilized (0.2 μm), transferred to Extetainer vials leaving no headspace, and stored at 4 °C for isotopic analysis of dissolved inorganic carbon (DIC). Finally, internal incubator gas was sampled for ambient and 2% CO_2_ experiments, as well as CO_2_ gas cylinders used to mix 2% CO_2_, following the sample collection protocol provided by the UC Davis Stable Isotope Facility.

The carbon isotope composition of bulk biomass (δ^13^C_biomass_), DIC (δ^13^C_DIC_), and CO_2_ (δ^13^C_CO2_) samples was determined at the UC Davis Stable Isotope Facility. δ^13^Cbiomass was analyzed using a PDZ Europa ANCA-GSL elemental analyzer interfaced to a PDZ Europa 20-20 isotope ratio mass spectrometer (Sercon Ltd.). δ^13^C_DIC_ (prepared by CO_2_ gas evolution using 85% phosphoric acid) and δ^13^C_CO2_ was measured by a GasBench II system interfaced to a Delta V Plus IRMS (Thermo Scientific). All carbon isotopic composition values are reported relative to the Vienna PeeDee Belemnite standard (V-PDB):

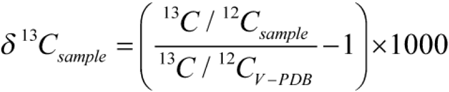

The carbon isotope fractionation associated with photosynthetic CO_2_ fixation (ε_p_) was calculated from measured δ^13^C_biomass_ and δ^13^C_CO2_ according to Freeman and Hayes (1992):

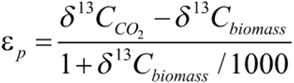

## Statistical analyses

Results for experimental analyses were presented as the mean and the sample standard deviation (SD) values of at least three independent experiments. Statistical significance was analyzed with the two-tailed *t*-test. The unpaired sample *t*-test assuming equal variances was used to compare the values obtained for different cyanobacterial strains and the paired sample *t*-test was used to compare the values for the same strain at different experimental conditions.

## Notes

### Competing Interest Statement

The authors have declared no competing interest.

### Summary of Updates

Additional analyses, updated text.

