## Supplementary Information for "Effects of CO_2_ and RuBisCO concentration on cyanobacterial growth and carbon isotope fractionation"

### *S. elongatus* culture carbon isotope dynamics – Extended results and discussion

Following dissolution of gaseous $CO_2$ ($CO_{2(g)}$) into the growth media, aqueous $CO_2$ ($CO_{2(aq)}$) is hydrated to other dissolved inorganic carbon (DIC) species, bicarbonate ($HCO_3^-$) and carbonate ($CO_3^{2-}$) (**Fig. S2**) and there are fractionations at each of these steps. At equilibrium, the isotopic compositions of DIC species can be determined by measurement of two parameters, such as the concentration of DIC and pH. However, our experimental system in the air treatment is not at equilibrium, as illustrated by the measured concentration and isotopic composition of DIC, as well as pH (**Table S2 and S2**).

In air, we measured a pH increase from ~7 to ~11 between the time of inoculation and biomass collection, whereas the pH remained near neutral under 2% $CO_2$. In air, $\delta^{13}C_{DIC}$ measured at the end of the culture period was, on average, isotopically depleted in $^{13}C$ by ~4–7‰ relative to $\delta^{13}C_{CO2(g)}$ sampled from the internal incubator atmosphere and displayed a high degree of variability across replicates. By contrast, $\delta^{13}C_{DIC}$ under 2% $CO_2$ was enriched in $^{13}C$ by ~8‰ compared to the $CO_{2(g)}$, indicating near-fequilibrium fractionation.

Under the experimental ambient air conditions, cyanobacteria are carbon-limited (observed by slower growth rates), experiencing low DIC concentrations in the media. At the biomass harvest time point, the high pH of the media (~11) would result in over 99% of DIC as carbonate ($CO_3^{2-}$), which rapidly interchanges with bicarbonate ($HCO_3^-$). The $HCO_3^-$ concentration is low due to its rapid cellular uptake as well as the high pH, driving a nearly unidirectional hydration of $CO_2$ to $HCO_3^-$, favoring $^{12}C$ with a kinetic isotope effect of ~18‰ (Yumol, Uchikawa, & Zeebe, 2020). A smaller, ~2‰ fractionation also exists between gaseous $CO_2$ and aqueous $CO_2$ (Zhang, Quay, & Wilbur, 1995). This is different from the 2% $CO_2$ treatment where we observe an equilibrium isotope effect. In this non-equilibrium system, the $\delta^{13}C$ of individual DIC species cannot be calculated based on equilibration equations and each species would have a $\delta^{13}C$ in between their kinetic and equilibrium isotope effects. Cyanobacteria can actively take up both aqueous $CO_2$ and bicarbonate from the media, and so $\delta^{13}C_{biomass}$ (-17.6 ± 0.3‰ and -17.9 ± 0.5‰ for WT and Syn02, respectively) may fall between the aqueous $CO_2$ and bicarbonate values (**Fig. S2**). Due to the carbon limitation of the experimental system in ambient air, we are unable to discern KIE differences between the RuBisCO enzymes of WT and Syn02 strains.

**Table S1. Growth parameters of *S. elongatus* strains, grown in unsparged cultures under varying $CO_2$ concentrations[a].**

| Strain | Atmosphere | Growth rate ($h^{-1}$) | Doubling time (h) | Carrying capacity ($OD_{750}$) |
|---|---|---|---|---|
| WT | Air | 0.018 ± 0.001 | 40 ± 2 | 2.85 ± 0.06 |
| | 2% $CO_2$ | 0.039 ± 0.001 | 18.0 ± 0.5 | 6.9 ± 0.2 |
| | 5% $CO_2$ | 0.039 ± 0.001 | 17.7 ± 0.4 | 5.52 ± 0.06 |
| Syn02 | Air | 0.018 ± 0.001 | 38 ± 1 | 2.8 ± 0.1 |
| | 2% $CO_2$ | 0.039 ± 0.001 | 17.6 ± 0.2 | 7.0 ± 0.2 |
| | 5% $CO_2$ | 0.044 ± 0.001** | 15.8 ± 0.6** | 5.6 ± 0.1 |

[a]Values are means of three biological replicates ± 1 SD. Asterisks indicate *t*-test result from comparison with WT for the same atmospheric condition. ** – $p$ <0.01.

**Table S2. $^{13}C/^{12}C$ isotopic and pH analysis of *S. elongatus* cultures under ambient air and 2% $CO_2$ (n = 7 for air values and n = 4 for 2% $CO_2$ values, except where noted).**

| Sample | Atmosphere | pH, start | pH, end | [DIC] (mM) | $\delta^{13}C_{biomass}$ (‰) | $\delta^{13}C_{DIC}$ (‰) |
|---|---|---|---|---|---|---|
| WT | Air | 6.98 ± 0.03 | 11.3 ± 0.2 | 0.22 ± 0.01 | -17.6 ± 0.3 | -20 ± 5 |
| | 2% $CO_2$ | dnr[a] | 7.26 ± 0.06 | 3.1 ± 0.3 | -57.20 ± 0.07 | -27.2 ± 0.3 |
| Syn02 | Air | 7.02 ± 0.01 | 11.2 ± 0.2 | 0.19 ± 0.03 | -17.9 ± 0.5 | -17 ± 8 |
| | 2% $CO_2$ | 6.10 ± 0.02 | 7.5 ± 0.1 | 3.9 ± 1.6 | -57.1 ± 0.4 | -26.6 ± 0.3 |
| BG-11 medium | Air | 7.02 ± 0.01 | 7.05 ± 0.02 | 0.160 ± 0.005 | n/a | -0.4 ± 2.5 |
| | 2% $CO_2$ | 6.09[b] | 6.4[b] | 0.65[b] | n/a | -29.3[b] |

[a]Did not record
[b]n = 1

**Table S3. $^{13}C/^{12}C$ isotopic analysis of *S. elongatus* culture headspace and $CO_2$ gas source.**

| Sample | $\delta^{13}C_{CO2}$ (‰) |
|---|---|
| Air, n = 3 | -12.91 ± 0.09 |
| 2% $CO_2$, n = 1 | -34.8 |
| $CO_2$ gas cylinder, n = 6 | -35.74 ± 0.08 |

**Table S4. Primers used in this study.**

| Primer[a] | Sequence (5'-3') | Description |
|---|---|---|
| F01 | GGCCAATAACCCAGGGATTTT GGAAAAAGCACTGTAATTC | Amplification of the *rbc* operon from the *S. elongatus* PCC 7942 gDNA with 20-nucleotide sequences that overlap with pAM4937 backbone to create the plasmid pSyn02. |
| R01 | GCCGGGGAGCTCCTTCATTTT CAGACTGCGTGGATAAG | |
| F02 | AGCGTCAGACCCCGTAGAAA AG | Amplification of the pAM4951 fragment that contains *ori* and *bom* from *E. coli* to construct the plasmid pSyn-01. |
| R02 | CCGCGGAGCTTGTCTGTAAG | |
| F03 | TAGCAGGCTTGGCGCGCCAA AAAAAAGC | Amplification of the pAM4951 fragment that contains the *aadA* gene to construct the plasmid pSyn-01. |
| R03 | AGGGAGAGACGACCCCTGGG TTATTGGCCGAC | |
| F04 | TAACCCAGGGGTCGTCTCTCC CTAGAGATATG | Amplification of the S. elongatus PCC 7942 gDNA fragment upstream of the rbc operon (contains the rbc promoter) with 20-nucleotide sequences that overlap with pAM4951 backbone to create the plasmid pSyn-01. |
| R04 | TTTCTACGGGGTCTGACGCTA TGCATCTACCGCCCCTAG | |
| F05 | CTTACAGACAAGCTCCGCGGG AACGCAGCCACAGGCCA | Amplification of the S. elongatus PCC 7942 gDNA fragment downstream of the rbc operon (contains the purK 3'UTR) with 20-nucleotide sequences that overlap with pAM4951 backbone to create the plasmid pSyn-01. |
| R05 | TTGGCGCGCCAAGCCTGCTAG GATGACCTCAG | |
| F06 | GACAATCCTGTTCTCCGGCA | Genotyping *S. elongatus* strains to screen for the *rbc* operon insert at NS2 (PCR product size: with *rbc* insert – 8,617 bp, without *rbc* insert – 1,970 bp). |
| R06 | ATCAACGCCGTACCCGTATC | |
| F07 | GGAGTCAATTCTGCAAGAGC | Genotyping *S. elongatus* strains to confirm the presence or absence of the *rbc* operon at the native site (PCR product size: with the *rbc* operon – 5,140 bp, without the *rbc* operon – 3,342 bp) R07 was also used to sequence the native *rbc* operon deletion site. |
| R07 | TCAAGCTCGGTCTACTGC | |
| F08 | GAATGCTCCGCTGGACTTGC | Sequencing the *rbc* operon insertion site at NS2. F08 was also used to sequence the native rbc operon deletion site. |
| R08 | TGTACTCGATTTGTGCAGCG | |
| F09 | ACCACCTTGGCAAAATGGTG | qPCR analysis of the expression of *rbcL* (ID: Synpcc7942_1426) that encodes the RuBisCO large subunit. |
| R09 | TTTGTCGCCTTCCAGTTTGC | |
| F10 | ATGTCGCTGCACGTTCAAAC | qPCR analysis of the expression of *purK* (ID: Synpcc7942_1428) that encodes N5- |
| R10 | TTGAGCCAATTTCGCAGTCG | |

| | | |
|---|---|---|
| | | carboxyaminoimidazole ribonucleotide synthase. |
| F11 | ATTACCTGCGCGACAACATG | qPCR analysis of the expression of *secA* (reference gene, ID: Synpcc7942_0289) that encodes the preprotein translocase subunit SecA. |
| R11 | TGCCCGCATGTATTTTTCGC | |
| F12 | ACGATTCGCGACAAACAACG | qPCR analysis of the expression of *ppc* (reference gene, ID: Synpcc7942_2252) that encodes phosphoenolpyruvate carboxylase. |
| R12 | AACACCGTTGGCTTGAACTG | |

[a]F - forward, R – reverse

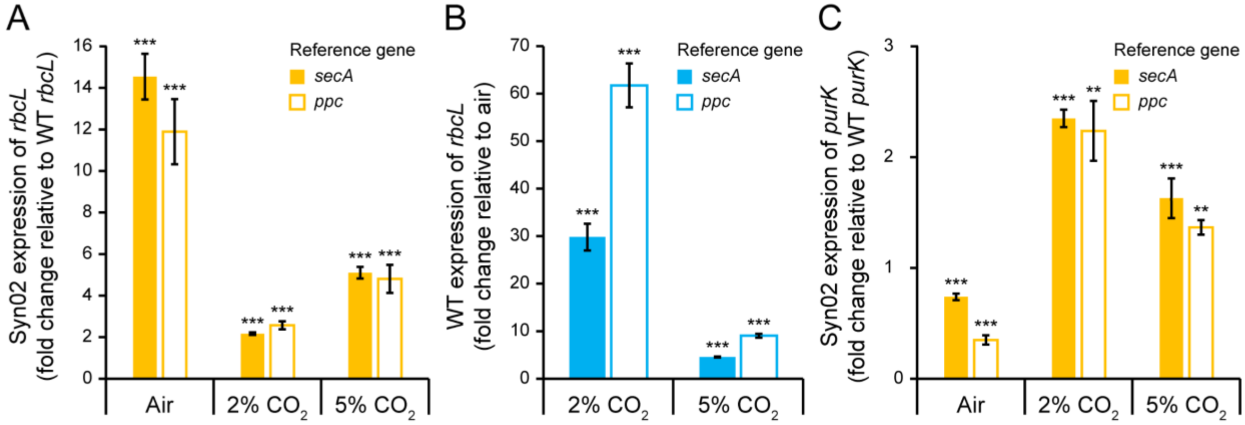


**Figure S1. Expression of RuBisCO during early growth phase of *S. elongatus* strains and expression of *purK*. A – Early growth phase expression of Syn02 *rbcL* relative to WT *rbcL*.** Cultures were collected at OD 0.5-1.5 for RT-qPCR analysis. Asterisks indicate *t*-test results compared to WT *rbcL* expression at the same growth condition. **B – Early growth phase expression of WT *rbcL* at 2-5% $CO_2$ relative to ambient air.** Asterisks indicate *t*-test results compared to WT *rbcL* expression in air. **C – Expression of Syn02 *purK* relative to WT *purK*.** Asterisks indicate *t*-test results compared to WT *rbcL* expression at the same growth condition. **A-C** – Data measured by RT-qPCR, normalized to reference genes *secA* and *ppc*. Columns represent mean fold expression for three biological replicates. Error bars on all graphs indicate 1 SD. ns – not significant; ** – $p < 0.01$; *** – $p < 0.001$.

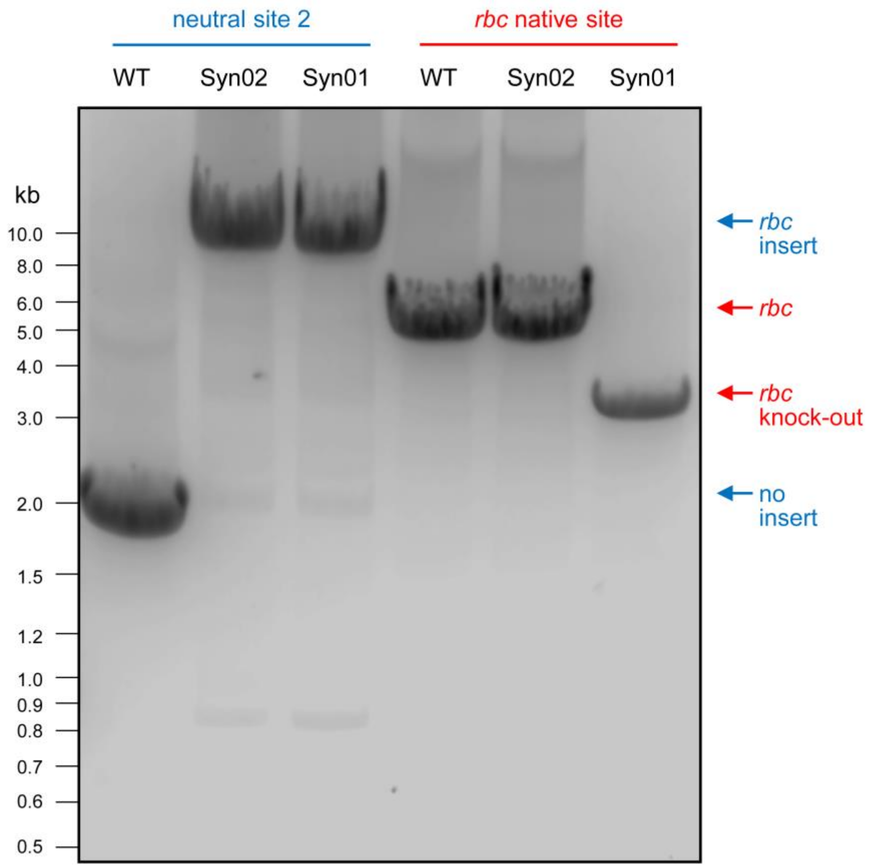


**Figure S2. Genotyping of *S. elongatus* strains by PCR.** Primers F06/R06 (**Table S4**) were used to confirm the *rbc* operon insertion at NS2 (*rbc* insert – 8,617 bp, no *rbc* insert – 1,970 bp). Primers F07/R07 were used to confirm the *rbc* operon presence at the native site (*rbc* present at its native site – 5,140 bp, *rbc* knocked out and replaced with the *aadA* gene – 3,342 bp).
